# Regulation of angiogenesis by signal sequence-derived peptides

**DOI:** 10.1101/2024.08.22.609269

**Authors:** Mean Ghim, Linyan Wei, Jae-Joon Jung, Erlinda The, Gunjan Kukreja, Afarin Neishabouri, Azmi A. Ahmad, M. Zawwad Raza, Arvene Golbazi, Keshvad Hedayatyanfard, Lei Nie, Jiasheng Zhang, Mehran M. Sadeghi

## Abstract

**Background:** The neuropilin-like, Discoidin, CUB and LCCL domain containing 2 (DCBLD2) is a transmembrane protein with an unusually long signal sequence (SS) composed of N-terminal (N) and C-terminal (C) subdomains, separated by a transition (tra) subdomain. DCBLD2 interacts with VEGFR-2 and regulates VEGF-induced endothelial cell signaling, proliferation and migration, as well as angiogenesis. The exact mechanisms by which DCBLD2 interacts with VEGFR2 to modulate VEGF signaling remain unclear.

**Methods:** Searching for VEGFR2 interacting DCBLD2 domains, we generated various constructs containing different DCBLD2 domain combinations and conducted co-immunoprecipitation and signaling studies in HEK 293T and endothelial cells. Several peptides were synthesized based on the identified domain, and their effect on VEGF signaling was assessed in vitro in cell culture and in vivo using matrigel plug and corneal micropocket assays. The effect of the lead peptide was further evaluated using a murine hindlimb ischemia model.

**Results:** DCBLD2 SS interacted with VEGFR2 and promoted VEGF signaling. SS was not cleaved in the mature DCBLD2 and its hydrophobic transmembrane ‘traC’ segment, but not the ‘N’ subdomain, was involved in DCBLD2-VEGFR2 interaction. The smallest unit in DCBLD2 SS that interacts with VEGFR2 was the L5VL5 sequence. Even after the central valine was removed, the L10 sequence mimicked the DCBLD2 SS traC’s effect on VEGF-signaling, while shorter or longer poly-leucine sequences were less effective. Finally, a synthetic traC peptide enhanced VEGF signaling in vitro, promoted VEGF-induced angiogenesis in vivo, and improved blood flow recovery following hindlimb ischemia.

**Conclusion:** DCBLD2 SS along with its derivative peptides can promote VEGFR2 signaling and angiogenesis. Synthetic peptides based on DCBLD2 SS hold promise as therapeutic agents for regulating angiogenesis. Importantly these findings refine the traditional view of signal sequences as mere targeting elements, revealing a role in cellular signaling. This opens new avenues for research and therapeutic strategies.

## INTRODUCTION

Targeting of newly synthesized proteins to the endoplasmic reticulum (ER) and eventually the plasma membrane of eucaryotic cells is mediated by signal sequences (SS). For type I transmembrane and secretory proteins, SS are typically 15 to 40 amino acid N-terminal extensions of the nascent proteins. During or after completion of ER translocation, many SS are cleaved by signal peptidase to generate a signal peptide (SP), while others remain an integral part of the mature protein, as signal anchor sequences ^1^. Generally, cleaved SPs are rapidly degraded. However, some SPs and their fragments are retained and have specific, post-targeting functions on their own ^2, 3^. To date, however, most known examples of such SS post-targeting functions are of viral origin ^4^.

A typical SP has a tripartite structure with a hydrophobic, α-helical core (*h* region) surrounded by an N-terminal polar *n* region, and a C-terminal *c* region that contains the signal peptidase cleavage site ^1^. Based on proteome analysis by machine-learning systems, it is suggested that exceptionally long SS may have a bipartite ‘NtraC’ domain structure with potentially functional N-terminal (N-domain, ‘N’) and C-terminal (C-domain, ‘C’) domains connected by a turn-rich linker area (transition area, ‘tra’) ^5^.

One such protein featuring an exceptionally long SS is Discoidin, CUB and LCCL domain containing 2 (DCBLD2), whose 66 amino acid SS is predicted to fit the ‘NtraC’ model ^6^. The gene encoding DCBLD2, also known as endothelial and smooth muscle cell–derived neuropilin-like protein (ESDN) or CLCP1, was initially cloned from human coronary artery ^7^ and highly metastatic lung cancer cells ^8^ as a protein with the longest known secretory signal sequence among eukaryotes. DCBLD2 is highly conserved among vertebrates and closely resembles the structure of neuropilins ^9, 10^, major co-receptors for vascular endothelial cell growth factor (VEGF) receptors (VEGFRs), which play a critical role in angiogenesis. DCBLD2 is upregulated in human, rat, and mouse remodeling arteries ^11, 12^, and regulates VEGF-induced endothelial cell (EC) proliferation, migration, and signal transduction, as well as developmental and adult angiogenesis ^13^.

Angiogenesis, the process of new blood vessel formation from existing vessels, plays a key role in a number of physiological processes, including development and growth, adult organ regeneration, and wound healing ^14, 15^. Angiogenesis is tightly controlled through numerous stimulators and inhibitors and its dysregulation, whether insufficient or excessive, contributes to a wide variety of diseases, including malignancy, ischemic heart and peripheral artery disease, neurodegeneration, and proliferative retinopathy ^16^. Accordingly, modulation of this process provides several therapeutic opportunities ^14^.

In this study, we identified the DCBLD2 SS, and more specifically its ‘traC’ segment, as a key domain of DCBLD2 that interacts with VEGFR2 and promotes VEGF/VEGFR2 signaling in EC. Furthermore, an exogenous synthetic FITC-traC peptide promoted angiogenesis in vivo. The smallest unit in DCBLD2 SS that interacted with VEGFR2 was the L5VL5 sequence. Even after the middle valine was removed, the L10 sequence could still mimic the effect of DCBLD2 SS traC on VEGF signaling. As such, in addition to establishing a novel post-targeting function of SS in regulating growth factor signaling, these findings introduce the DCBLD2 traC and its derivatives as promising candidates for therapeutic angiogenesis.

## MATERIALS AND METHODS

### Sex as a biological variable

In vivo studies exclusively used male mice to reduce variability. In vitro studies, performed with mixed populations of cells, suggest that the findings are sex independent.

### Cell culture

HEK 293T cells were maintained in Dulbecco’s Modified Eagle Medium (DMEM, Gibco) supplemented with 10% heat inactivated fetal bovine serum (FBS, Sigma Aldrich) and 1% penicillin/streptomycin (Gibco). For endothelial cell culture, primary human umbilical vein endothelial cells (HUVECs) were obtained from Yale Vascular Biology & Therapeutics Program Tissue Culture program and cultured on gelatin-coated cultureware with Medium 199 (Gibco) supplemented with 20% FBS, 1% endothelial cell growth supplement (ECGS), 2 mM L-glutamine, and 1% penicillin/streptomycin. MLEC were isolated from 4-week-old wild-type and *Dcbld2^-/-^*mice. Mice were anesthetized and lungs harvested, rinsed in Hank’s Balanced Salt Solution (HBSS, Sigma Aldrich), cut into small pieces, and subjected to enzymatic digestion with a 1mg/ml collagenase solution for 45 minutes. The tissue solution was then mechanically disrupted, passed through a 70 µm strainer, and cells were collected by centrifugation at 350g for 5 minutes and resuspended in complete Medium 199. CD31-positive cells were separated using immunomagnetic separation with CD31-conjugated beads, followed by detachment of the cells from the beads and cultured until 70-80% confluent. Finally, cells were subjected to another round of immunoselection with CD102-conjugated beads following the same procedure and detachments. The purity of isolated MLECs was verified by assessing the expression of CD31 with flow cytometry. Preparation of antibody conjugated magnetic beads was carried out according to manufacturer’s (Thermo Fisher) instructions. A 0.25% trypsin-EDTA (Gibco) solution was used together with standard lab protocols for passing adherent cells.

### Plasmid construction and transfection

The constructs used in this study are schematically shown in Fig. 1A and 1F. The genes encoding different protein fragments were cloned into a pCDNA3.0 plasmid between HindIII and KpnI restriction sites ^8^. All constructs were confirmed by PCR or DNA sequencing. Transfection of cells with plasmid DNA was carried out using Lipofectamine 3000 (Thermo Fisher) according to the manufacturer’s instructions. 24 hours after transfection, cells were washed and cultured in DMEM supplemented with 10% FBS for another 24 hours before use.

**Fig. 1.**
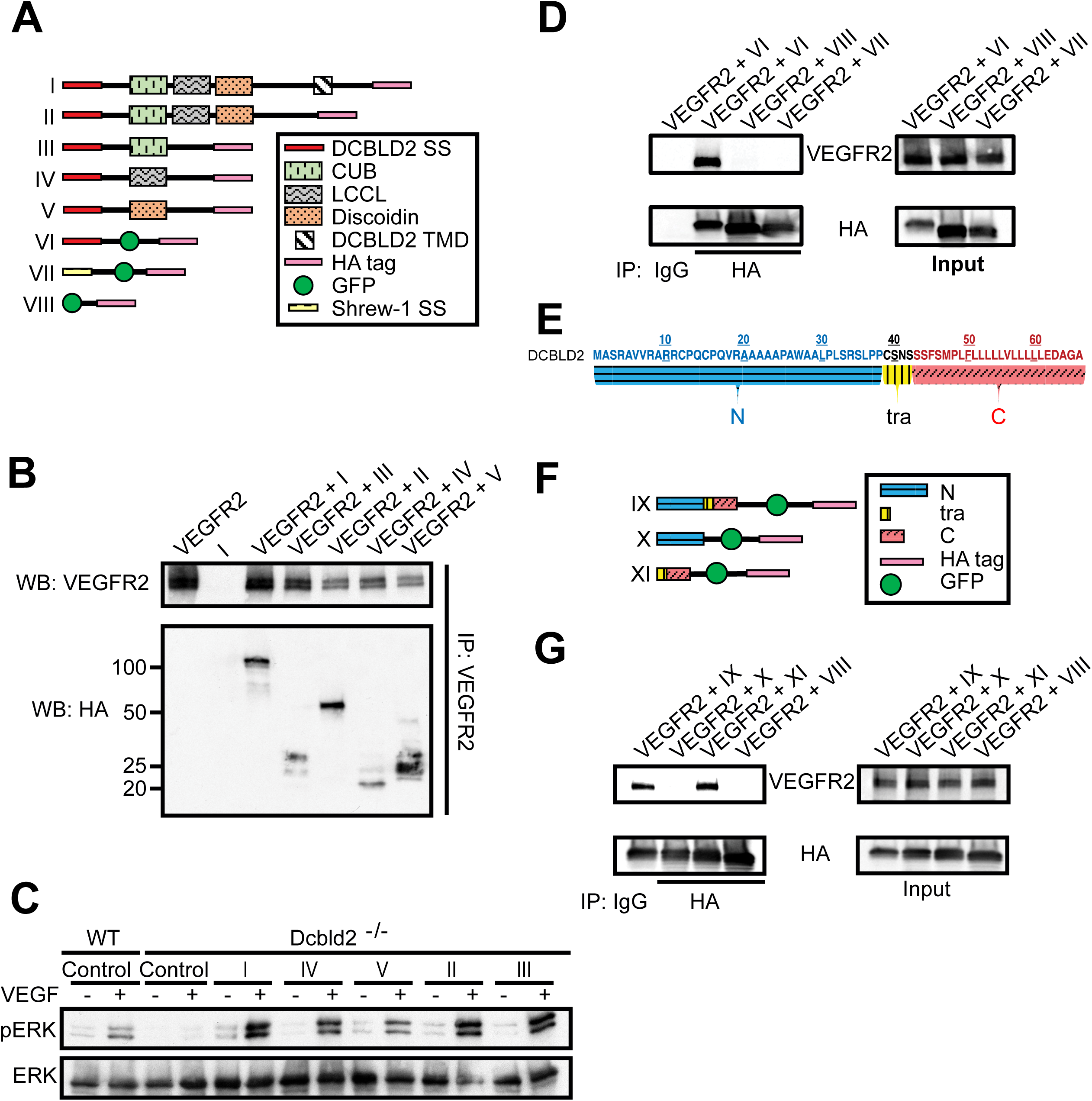
The DCBLD2 SS traC subdomain is involved in DCBLD2 interaction with VEGFR2. (A) Schematic representation of constructs I-VIII, each with various combinations of DCBLD2 domains tagged with HA (I-V), DCBLD2, or Shrew-1 SS fused with GFP and HA tags (VI, VII) or GFP-HA alone (VIII). (B) Co-immunoprecipitation of the products of constructs I-V with VEGFR2 following transfection of each construct with VEGFR2 (or transfection of construct I alone without VEGFR2) in HEK 293T cells. VEGFR2 immunoprecipitation is followed by Western blotting for HA and VEGFR2. (C) Western blot analysis of the effects of constructs I-V on VEGF-induced ERK phosphorylation following their transfection in *Dcbld2^-/-^* murine lung endothelial cells transfected. (D) Co-immunoprecipitation of the products of constructs VI-VIII with VEGFR2 following their transfection in HEK 293T cells. HA immunoprecipitation is followed by Western blotting for HA and VEGFR2. (E) NtraC model of DCBLD2 SS. (F) Schematic representation of constructs IX-XI encompassing different DCBLD2 SS subdomains fused with GFP and HA tags (IX-XI) or GFP-HA alone (VIII). (G) Co-immunoprecipitation of the products of constructs IX-XI with VEGFR2 following their transfection in HEK 293T cells. HA immunoprecipitation is followed by Western blotting for HA and VEGFR2. SS: signal sequence; TMD: transmembrane domain. HA: Hemagglutinin; GFP: green fluorescent protein; IP: immunoprecipitation; WB: western blot; WT: wild type.

### Immunoblotting and immunoprecipitation

Cells were lysed on ice using ice-cold RIPA buffer (Thermo Fisher) supplemented with protease and phosphatase inhibitor cocktails (Roche). The lysates were centrifuged at 14,000g for 15 minutes at 4°C, and supernatant was collected. Protein concentrations were determined using BCA assays (Thermo Fisher). Equal amounts of protein were subjected to SDS-PAGE and transferred onto 0.2µm PVDF membranes (Bio-Rad). Membranes were blocked for 1 hour with a 5% BSA solution (Sigma Aldrich) in TBST (Cell Signaling). For immunoblotting, membranes were incubated overnight with primary antibodies at 4°C and for 1 hour at room temperature with secondary antibodies. The blots were developed using an ECL substrate (Thermo Fisher).

For co-immunoprecipitation studies, Dynabeads™ Protein A Immunoprecipitation Kit (Invitrogen) was used. All steps were carried out according to the manufacturer’s protocol. After completing the protocol, beads were removed with a magnet and the resultant supernatant was collected for analysis.

### Immunofluorescence staining

Cell monolayers and OCT embedded tissue sections were fixed with 4% paraformaldehyde (PFA) for 15 minutes and blocked for 1 hour with 5% BSA at room temperature. Cells and tissue sections were incubated overnight at 4 °C with primary antibodies followed by incubation with secondary antibodies) for 1 hour at room temperature. Primary and secondary antibody dilutions were made with 5% BSA, based on the manufacturers’ recommended concentrations for each antibody. Cell nuclei were stained with DAPI (Thermo Fisher) and coverslips were secured with mounting media (Thermo Fisher) and allowed to cure for 48 hours Images were acquired with inverted Leica SP8 confocal or DMi8 widefield microscopes.

### Synthetic peptides

Thermo Fisher Scientific was commissioned to produce the synthetic traC peptide using the following sequence: [FITC][Ahx]CSNSSSFSMPLFLLLLLVLLLLLEDAGA[COOH]. Mass spectrum and HPLC data reported a molecular mass of 3483.18 Da and purity of >98%. Due to the hydrophobic properties of the peptide, traC was reconstituted in DMSO, aliquoted to avoid freeze-thaw cycles, and stored at -80°C. For experiments, aliquots were diluted and vortexed before application, ensuring even distribution in aqueous media and preventing areas of localized concentration during treatment. FITC-L10 and FITC-scramble peptide, with a sequence of [FITC][Ahx]LLLLLLLLLLEDAGA[COOH] and [FITC][Ahx]LSLESALMLFLCLDLSGVFLSLPLSNL[COOH] respectively, were obtained from Thermo Fisher Scientific. Mass spectrum and HPLC data reported a molecular mass of 2094.58 Da and 3412.10 Da for FITC-L10 and FITC-scramble respectively and a purity of >98% for both. Antennapedia peptide was obtained from GeneScript, with mass spectrometry and HPLC analysis reporting a molecular mass of 2749.28 Da and a purity of 96.6%.

### Matrigel plug angiogenesis assay

One µmol of traC, with or without 2 μg/ml VEGF, was mixed into 300 µl of growth factor-reduced Matrigel (Corning) at 4℃. The mixture was subcutaneously injected into the opposite iliac regions of 8–10-week-old C57BL/6J WT mice. Five days post-injection, mice were euthanized and Matrigel plugs were collected and embedded in OCT. The plugs were frozen, sectioned, and stained with goat polyclonal anti-CD31 (R&D Systems) overnight at 4°C, followed by Alexa Fluor 594-labeled chicken anti-goat IgG (Thermo Fisher) for 1 hour at room temperature. Sections were imaged using a widefield microscope (Leica DMi8). When imaging, areas containing the highest CD31 staining at the edge of the sections were imaged and the percentage of the CD31-positive area relative to the total imaged area was quantified blindly and reported.

### Pellet preparation and corneal micro-pocket angiogenesis assay

Pellets were prepared 24 hours before implantation. Initially, 5 µl of a 12% (w/v) solution of poly(2-hydroxyethyl methacrylate) (poly-HEMA; Sigma Aldrich) dissolved in ethanol and 1 µl of a 10% (w/v) sucralfate solution (Cayman Chemical) dissolved in PBS were mixed.

Subsequently, either 4 µl of FITC-traC (0.75 µg/µl), 4 µl of recombinant human VEGF (1 µg/µl, R&D Systems), or 4 µl of a solution containing both FITC-traC (0.75 µg/µl) and VEGF (1 µg/µl) was added and mixed. Vehicle controls consisted of 4 µl of the diluted base solvent used to dissolve FITC-traC or VEGF. From the resulting 10 µl mixture, 0.2 µl of the solution was pipetted onto parafilm to create pellets and allowed to dry. The resulting pellets contained either 60 ng of FITC-traC, 80 ng of VEGF, a combination of both, or the vehicle controls. Implantation of the pellets was performed on 12-week-old C57BL/6J mice (Jackson Laboratory) as previously described ^17^. Briefly, using a micro-knife (Ambler Surgical), a 1 mm incision was made on the cornea, approximately 1 mm from the limbus, into which a pellet was inserted. Each mouse was implanted with two pellets, one under each cornea. 13 days post implantation, corneas were harvested and fixed in 4% paraformaldehyde on ice for 1 hour, followed by permeabilization and blocking with a 5% BSA solution containing 0.1% Triton X-100 (Sigma Aldrich). Goat polyclonal anti-CD31 (R&D systems) was prepared in blocking buffer and incubated with the corneas overnight, with gentle agitation at 4℃. For secondary antibody labeling, corneas were incubated with Alexa Fluor 594-labeled chicken anti-goat IgG (Thermo Fisher) for 2 hours at room temperature. Finally, corneas were flattened, whole-mounted onto glass slides, and imaged by confocal microscopy using the tile scan function with a z-stack; maximum projections were used for the quantification. Images were thresholded to distinguish CD31-positive staining from the background. To reduce sampling bias, the entire flat-mounted cornea was analyzed blindly. The total cornea area was defined with the innermost vessel of the limbus arcade as the border. The CD31 positive area was quantified and reported as a percentage of the ROI.

### Hindlimb ischemia

For Hindlimb Ischemia, 12-week-old C57BL/6J mice (Jackson Laboratory) were subjected to femoral artery ligation and analyzed by Laser Doppler imaging to measure blood flow in the hindlimb as previously described with minor modifications ^13^. Briefly, the femoral artery was ligated proximal to the superficial epigastric artery branch and all branches between the two ligatures were ligated. The arterial segment between the ligatures was then excised.

Osmotic mini pumps (Alzet) were primed and prepared to deliver 1.74mg/day of traC for 7 days to the ligated limb based on a pumping rate of 1.05ml/hr, according to the manufacturer’s instructions. Following ligation, the osmotic mini pump was subcutaneously inserted into the back of the mouse, and a catheter (Alzet) connected to the pump was guided and secured adjacent to the ligated thigh. The animals were randomly assigned to treatment or control groups. Laser Doppler flow images of the foot were acquired before, immediately after surgery, and on days 3, 5, and 7 post-surgery. Images were recorded blinded to treatments with an infrared laser doppler imager (LDI; Moor Instruments) at 37.0-37.5 °C under anesthesia with ketamine/xylazine (100/10 mg/kg). The data were analyzed with the Moor LDI image processing software V5.00 and the results were reported as a ratio between the ischemic and non-ischemic foot.

After 7 days of treatment, the mice were euthanized and the thigh and the leg were collected in OCT, frozen, and sectioned. Four sections, approximately 2 mm spaced apart, from each thigh were stained for CD31. The percentage of the CD31-positive area was calculated for each of the four sections relative to the total area of each thigh and the results were averaged to obtain a representative percentage of the CD31-positive area within each thigh. Quantification of *cdh5* gene expression was performed using RT-PCR with *Gapdh* used as the housekeeping gene. Results were reported as 2^-ΔCt^. Fifty sections, each 5 µm thick, were collected at 8 locations along the thigh or the leg at 1 mm intervals and pooled for each animal. Total mRNA was isolated using the RNeasy kit (Qiagen), and reverse transcription (RT) was carried out with the QuantiTect RT kit (Qiagen). Real-time PCR was conducted in triplicates using cDNA with TaqMan gene expression assays, following the manufacturer’s instructions. Tissue analysis was performed blinded to treatment status.

### Statistical analysis

Results are expressed as the mean ± standard deviation. All statistical analyses were performed using GraphPad Prism 10. Data were checked for normality with the Shapiro-Wilk test. For data that did not pass the Shapiro-Wilk test, non-parametric tests were used. Statistical significance was determined by a two-way analysis of variance (ANOVA, followed by Tukey’s post hoc test, for multiple groups, and a student t-test or a Mann-Whitney U test for comparison between two groups. A p-value<0.05 was considered significant.

### Study approval

All studies were performed under protocols approved by Yale University and Veterans Affairs Connecticut Healthcare System Institutional Animal Care and Use and Human Investigational Committees.

## RESULTS

### The DCBLD2 SS traC subdomain is involved in DCBLD2 interaction with VEGFR2

Distinct functional and structural units of DCBLD2 include an unusually long SS, which in tandem with CUB, LCCL, and Discoidin domains form the extracellular segment. A transmembrane domain and an intracellular segment comprise the rest of the molecule. To identify the key domain(s) of DCBLD2 that interact with VEGFR2, we generated several plasmids expressing various combinations of different DCBLD2 domains linked to a C-terminal hemagglutinin (HA) tag (Fig. 1A I-V. Upon co-transfection with VEGFR2 in HEK 293T cells each of these constructs expressed the appropriate-size protein (Fig. S1). Surprisingly, co-immunoprecipitation with an anti-VEGFR2 antibody showed that all these products complex with VEGFR2 (Fig. 1B). Furthermore, while as reported previously ^13^, VEGF signaling was diminished in *Dcbld2*-deficient murine lung endothelial cells (MLEC), all these DCBLD2 constructs enhanced VEGF-induced ERK phosphorylation upon transfection in *Dcbld2*^-/-^ MLEC (Fig. 1C). This led us to speculate that a common component of the constructs, i.e., the SS, might be involved in its interaction with VEGFR2 and enhancing VEGFR2 signaling.

To test this possibility, and to rule out HA as the VEGFR2 interacting component, we constructed additional plasmids expressing DCBLD2 SS, Shrew-1 SS (another unusually long SS with a similar ‘NtraC’ domain structure ^5^) or no SS in tandem with C-terminal green fluorescent protein (GFP)-HA tags (Fig. 1A VI-VIII). Upon co-transfection of these constructs with VEGFR2 in HEK 293T cells and HA immunoprecipitation, VEGFR2 could be pulled down by GFP-HA-tagged DCBLD2 SS, but not Shrew-1 SS or GFP-HA without any SS (Fig. 1D). In addition, despite a similar predicted size, the product of DCBLD2 SS-GFP-HA plasmid appeared larger on western blots than the products of Shrew-1 SS-GFP-HA or GFP-HA constructs (Fig. 1D). This raised the possibility that the Shrew-1 SS but not the DCBLD2 SS is cleaved in the final product. Indeed, the SignalP 6.0 server, which can predict the location of the signal peptide cleavage site ^18^, suggested that unlike Shrew-1, the N-terminal SS is not cleaved in DCBLD2 (Fig. S2). To confirm this experimentally, we transfected HEK 293T cells with a dually tagged DCBLD2 plasmid with N-terminal FLAG and C-terminal HA tags. Immunoprecipitation of the product with an anti-HA antibody followed by FLAG and DCBLD2 immunoblotting showed a single (non-cleaved) product (Fig. 3). Replacing one or both alanine residues in the putative A-X-A signal peptidase cleavage site ^19^ of DCBLD2 with glycine led to similarly sized products, indicating that the DCBLD2 SS remains uncleaved (Fig S3). Finally, VEGF-induced ERK phosphorylation in *Dcbld2^-/-^* MLEC was enhanced upon transfection with the DCBLD2 SS-GFP-HA construct (Fig. S4). Taken together, these data suggest the DCBLD2 SS remains an integral component of the mature DCBLD2 and is a key domain in the DCBLD2 interaction with VEGFR2 that modulates downstream VEGF signaling.

Based on the ‘NtraC’ model, the DCBLD2 SS ’N’ subdomain contains 38 amino acid residues, and the 24 amino acid C-terminal residues constitute the ‘C’ subdomain, with those two separated by a short ‘tra’ subdomain (Fig. 1E). Hydrophobicity analysis by ProtScale ^20^ (Fig. S5) and the transmembrane helix prediction by DeepTMHMM^21^ (Fig. S6) indicated significant differences between different DCBLD2 SS subdomains, with the ‘C’ subdomain adjacent to ‘tra’ predicted to be hydrophobic and transmembrane. To identify the subdomain of DCBLD2 SS that interacts with VEGFR2, we synthesized additional constructs IX-XI expressing different DCBLD2 SS subdomains (Fig. 1F), which were co-transfected with VEGFR2 into HEK 293T cells. Co-immunoprecipitation studies indicated that the ‘traC’ subdomain, but not the ‘N’ subdomain, is involved in the DCBLD2-VEGFR2 interaction (Fig. 1G).

### FITC-TraC enhances VEGF/VEGFR2 signaling in EC

Given the interaction between the DCBLD2 SS traC fragment and VEGFR2, we sought to investigate whether a synthetic traC peptide, labeled with FITC for tracking purposes, can modulate VEGF signaling in EC. As a first step in evaluating the effects of FITC-traC in EC, we followed by fluorescence imaging the fate of the labeled peptide added to EC culture media.

Interestingly, FITC-traC was rapidly taken up by HUVEC with the peptide detected in the cytoplasm as early as 1 minute after its addition to the medium. The cytoplasmic accumulation of the peptide increased over time and by 15 minutes it was also detectable in the cell membrane (Fig. 2A), where VEGFR2 is also located (Fig. 2B). Accordingly, the 15-minute time point was selected for the next set of studies.

**Fig. 2.**
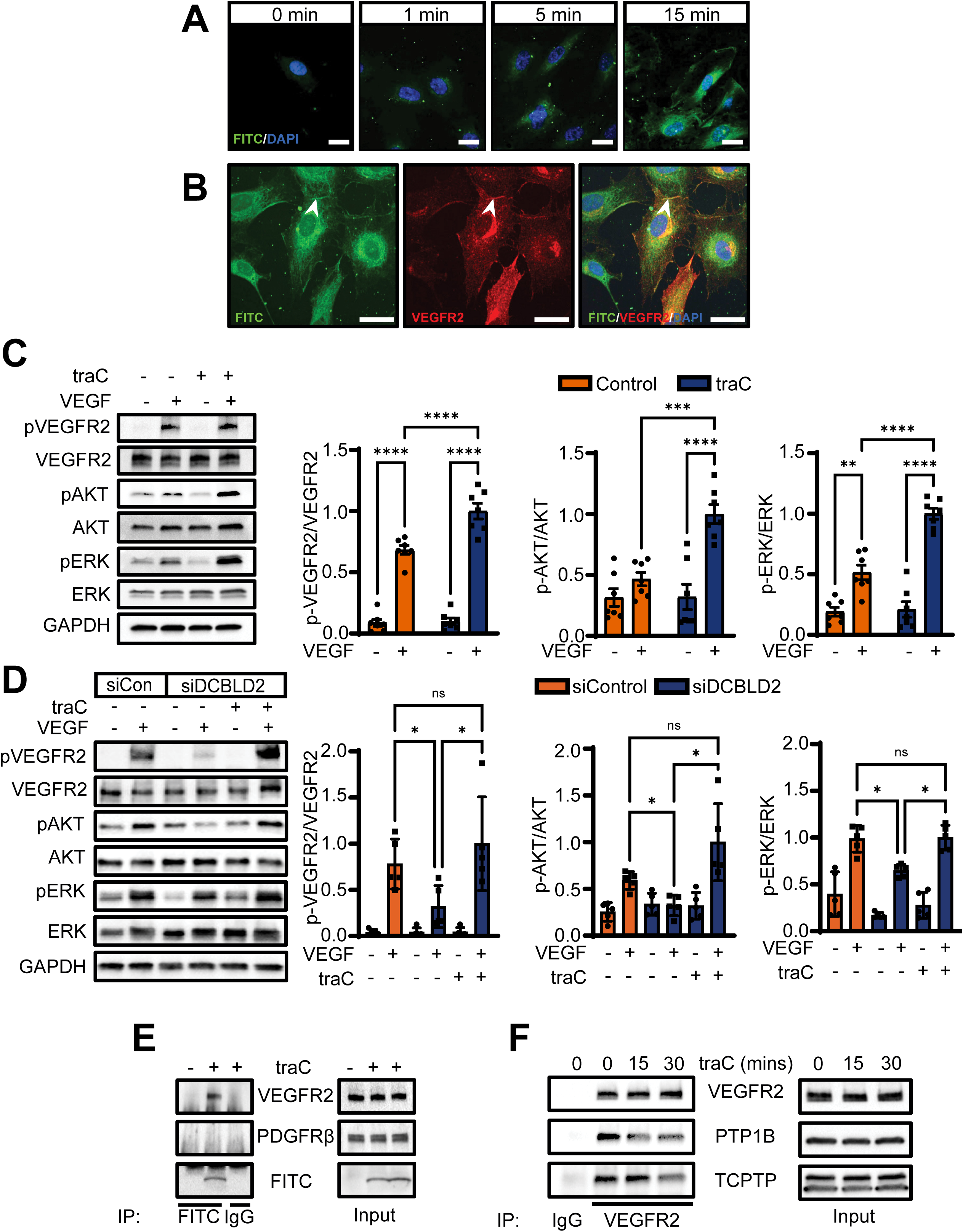
FITC-traC enhances VEGF/VEGFR2 signaling in endothelial cells. (A) Illustrative immunofluorescence images showing FITC-traC (green) peptide penetration into the cells over time. Nuclei are stained blue with DAPI. Bar = 20μm. (B) Illustrative immunofluorescent images showing co-localization of FITC-traC (green) with VEGFR2 (red) on the cell membrane (arrowheads) 15 minutes after adding FITC-traC to endothelial cells. Nuclei are stained blue with DAPI. Bar = 20μm. (C) Western blot analysis of VEGF induced VEGFR2, ERK, and Akt phosphorylation with and without FITC-traC (n=7). (D) Western blot analysis of the effect of FITC-traC on VEGF induced VEGFR2, ERK, and Akt phosphorylation in endothelial cells following siRNA-mediated DCBLD2 downregulation (n=5). (E) Co-immunoprecipitation of FITC-traC with VEGFR2 but not PDGFRβ following the addition of FITC-traC to endothelial cell cultures. FITC immunoprecipitation is followed by Western blotting for VEGFR2, PDGFRβ, or FITC. (F) Effect of FITC-traC on protein tyrosine phosphatases PTP1B and TCPTP interaction with VEGFR2 in endothelial cells. VEGFR2 immunoprecipitation is followed by Western blotting for VEGFR2, PTP1B or TCPTP. PTP1B: Protein tyrosine phosphatase 1B; TCPTP: T cell protein tyrosine phosphatase. ns: not significant; *: P<0.05, **: P<0.01, ***: P<0.001, ****: P<0.0001. Statistical significance was determined by two-way ANOVA and Tukey’s multiple comparison post hoc test (C) or by repeated measures one-way ANOVA and Holm-Šídák’s multiple comparisons post hoc test (D).

Next, we evaluated the effect of synthetic DCBLD2 FITC-traC on VEGF signaling. Pretreatment of HUVEC with FITC-traC peptide (1μM in 1% DMSO, 15 minutes), but not a scrambled homologue, significantly increased VEGF-induced VEGFR2, ERK, and AKT phosphorylation compared to HUVEC pre-treated with vehicle control (Fig. 2C and Fig. S7). A dose-response study showed that the effect of FITC-traC peptide on VEGF signaling is concentration-dependent (Fig. S8). Pre-heating of FITC-traC peptide (120 °C, 50 minutes) abrogated its effect on VEGF signaling (Fig. S9). To explore the molecular mechanisms of this observation, we first determined whether the effect of FITC-traC is simply related to its cell penetrating property. A similar set of experiments showed that unlike FITC-traC, pre-treatment of HUVEC with a classical cell penetrating peptide, antennapedia, does not affect downstream VEGF signaling (Fig S10). Next, we sought to determine whether FITC-traC peptide can reverse the inhibitory effect of DCBLD2 downregulation on VEGF signaling in EC. As expected ^13^, DCBLD2 siRNA significantly reduced VEGF-induced VEGFR2 and ERK phosphorylation in HUVEC. This impaired VEGF/VEGFR2 signaling was fully restored upon pre-treatment of cells with FITC-traC peptide (Fig. 2D). Importantly, VEGFR2, but not PDGFRβ, could be co-immunoprecipitated with FITC-traC, when HUVEC were pretreated with the peptide for 15 minutes (Fig. 2E), indicating that like endogenous DCBLD2, the exogenous peptide interacts with VEGFR2. The specificity of VEGFR2 interaction with FITC-traC was confirmed following co-transfection of different tyrosine kinase receptors in HEK 293 cells (Fig. S11). As the effect of DCBLD2 in regulating VEGF signaling has been linked to regulation of VEGFR2-protein tyrosine phosphatase interaction, we evaluated the effect of synthetic FITC-traC peptide on VEGFR2 interactions with protein tyrosine phosphatase 1B (PTP1B) and T cell protein tyrosine phosphatase (TCPTP) ^13^. Addition of FITC-traC peptide to HUVEC markedly reduced PTP1B and TCPTP co-precipitation with VEGFR2 (Fig. 2F), suggesting that the effect of FITC-traC on VEGF signaling parallels, at least in part, the effect of endogenous DCBLD2 on VEGF signaling.

### DCBLD2 SS FITC-traC peptide promotes VEGF and ischemia-induced angiogenesis in vivo

The effect of DCBLD2 SS FITC-traC peptide on in vivo response to exogenous VEGF was evaluated in matrigel plug and cornea micropocket angiogenesis assays. Visual inspection of matrigel plugs collected 5 days after implantation showed apparently more neovascularization in the presence of FITC-traC and VEGF, compared to VEGF alone (Fig. 3A). This was confirmed by immunostaining which showed significantly more CD31 in matrigel plugs containing FITC- traC and VEGF compared to VEGF alone, while in the absence of VEGF, there was no difference between FITC-traC and control matrigel plugs (Fig. 3B). Similarly, in the cornea micropocket assay, FITC-traC enhanced angiogenesis in the presence of VEGF. This difference was readily detectable on visual inspection of corneas and was confirmed by CD31 immunostaining (Fig. 3C-D).

**Fig. 3.**
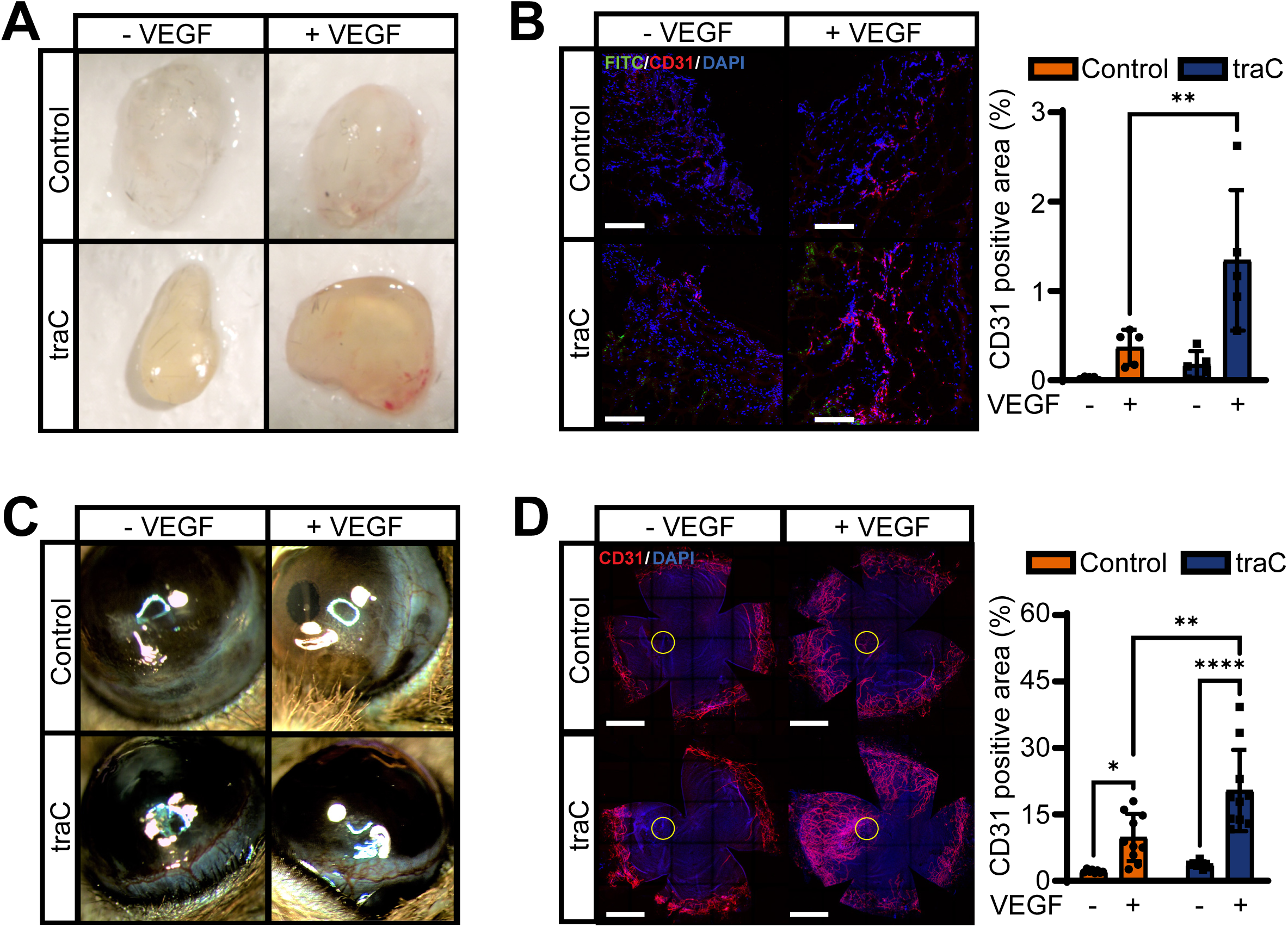
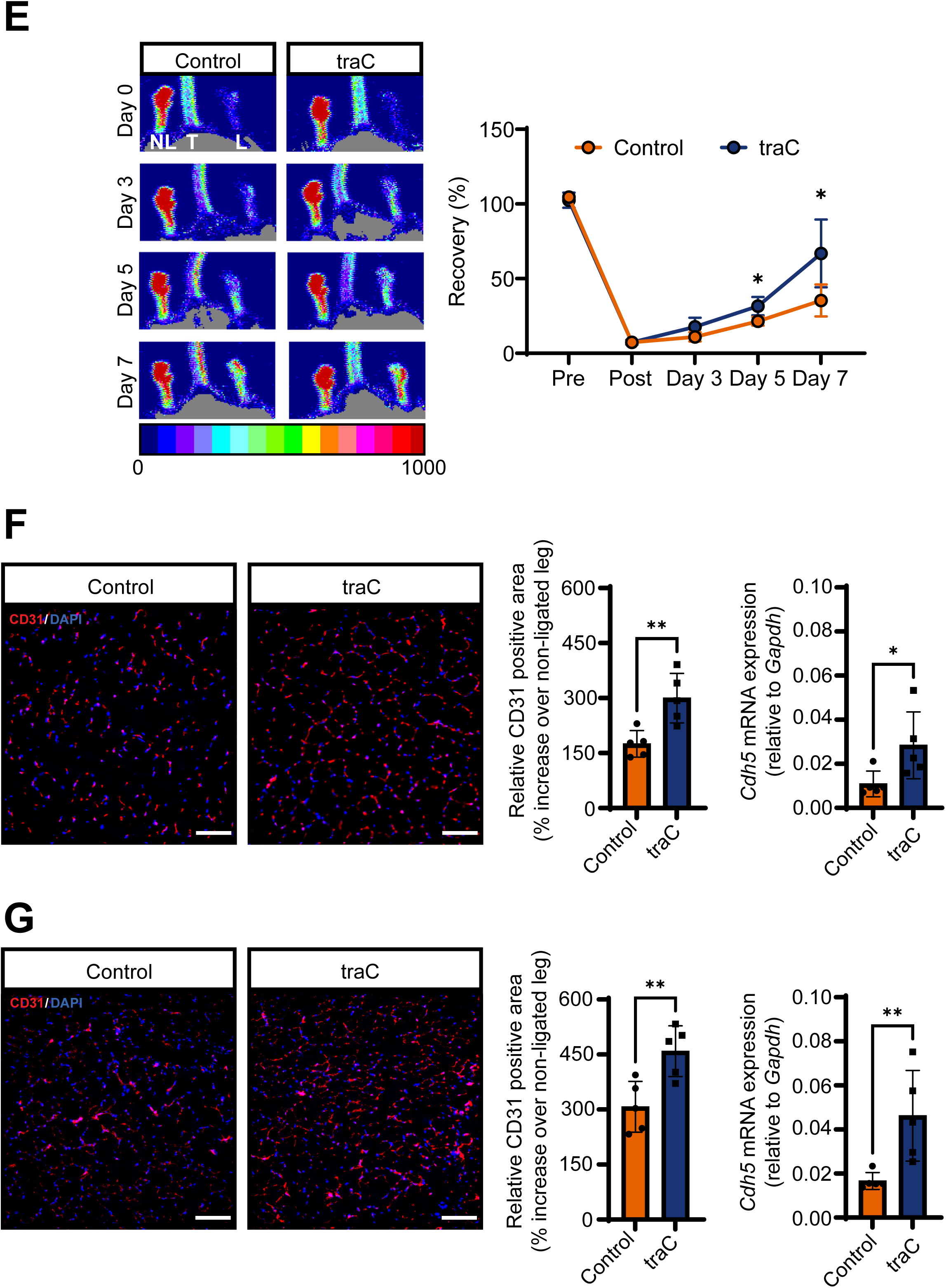
The DCBLD2 SS FITC-traC peptide promotes VEGF and ischemia-induced angiogenesis in vivo. (A, B) Illustrative photographs (A) and CD31 immunofluorescent staining (red, n=5, B) of matrigel plugs with or without VEGF and FITC-traC, collected after 5 days post-implantation. Nuclei are stained blue with DAPI. Bar = 200μm. (C, D) Illustrative slit-lamp photographs (C) and CD31 immunofluorescent staining (red, n=10, D) of corneas implanted with sustained-release pellets containing either FITC-traC or DMSO and with or without VEGF. Bar= 1mm. (E-G) Illustrative laser Doppler images and quantification of hindlimb blood flow recovery after femoral artery ligation in mice treated with FITC-traC compared to control animals. L: ligated, NL: non-ligated, T: tail. (n=5, E). Illustrative examples of the thigh (F) and calf (G) CD31 immunofluorescent staining (red) and its quantification, as well as *Gapdh-*normalized *Cdh5* expression by real time PCR at day 7 post femoral artery ligation in mice treated with FITC-traC compared to control animals (n=5). Nuclei are stained blue with DAPI. Bar=200μm. *: P<0.05, **: P<0.01. Statistical significance was assessed by two-way ANOVA and Tukey’s multiple comparison test (B and D) or Student t-test (E, F and G).

Next, we evaluated the effect of FITC-traC on blood flow recovery and angiogenesis following femoral artery ligation in mice. Evaluation of hindlimb blood flow by laser Doppler imaging showed improved blood flow recovery in animals continuously treated with FITC-traC, detectable as early as 5 days after femoral artery ligation, compared to control animals (Fig. 3E). Histological analysis of the hindlimb muscles collected 7 days after femoral artery ligation showed significantly higher thigh and calf muscle capillary density, as assessed by immunostaining (for CD31) and quantitative RT-PCR (for VE-cadherin) in FITC-traC treated mice relative to control animals at 7 days (Fig. 3F, G). As expected, no difference was found in the non-ligated hindlimb VE-cadherin expression between the two groups of animals (Fig. S12).

### A poly leucine decapeptide mimics the effect of DCBLD2 SS traC on VEGF signaling

As a prelude to future evaluation of related signal sequences in therapeutic applications, we sought to identify its smallest sequence that interacts with VEGFR2. The DCBLD2 SS traC was split into two parts with an N-terminal part 1 (CSNSSSFSMPLF), and a C-terminal part 2 (LLLLLVLLLLLEDAGA) encompassing the poly-leucine sequence. Following co-transfection of plasmids expressing VEGFR2 with GFP-HA-tagged part 1 or part 2 into HEK 293 cells, part 2, but not part 1 could be co-immunoprecipitated with VEGFR2 (Fig. 4A). Part 2 was then split into two sub-parts: LLLLLVLLLLL (L5VL5) and EDAGA, each connected to a C-terminal GFP-HA tag. Following co-transfection with VEGFR2 in HEK 293 cells, only the sequence L5VL5, and not EDAGA, could co-precipitate with VEGFR2, indicating that this segment is involved in the DCBLD2 SS interactions with VEGFR2 (Fig. 4B).

**Fig. 4.**
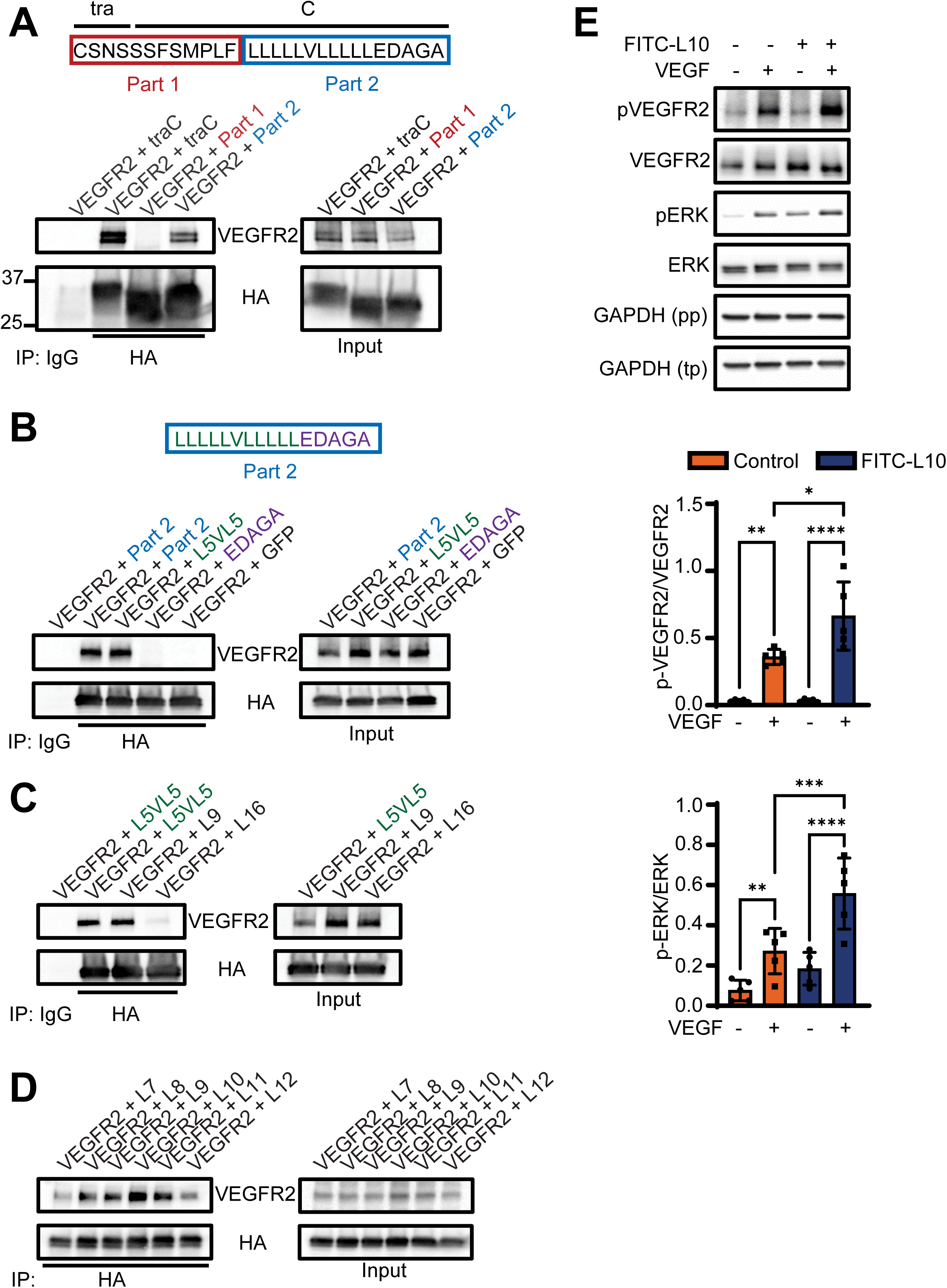
A poly leucine decapeptide mimics the effect of DCBLD2 SS traC on VEGF signaling. (A) Co-immunoprecipitation of the products of HA-tagged traC Part 1 or Part 2 constructs with VEGFR2 following their co-transfection in HEK 293T cells. HA immunoprecipitation is followed by western blotting for HA and VEGFR2. (B) Co-immunoprecipitation of the products of HA-tagged traC Part 2 or its derivative constructs or a control HA-tagged GFP with VEGFR2 following their co-transfection in HEK 293T cells. HA immunoprecipitation is followed by western blotting for HA and VEGFR2. (C) Co-immunoprecipitation of the products of HA- tagged traC Part 2 L5VL5 sequence, or 9 or 16 poly-L-leucine constructs with VEGFR2 following their co-transfection in HEK 293T cells. HA immunoprecipitation is followed by western blotting for HA and VEGFR2. (D) Co-immunoprecipitation of the products of HA- tagged polyleucine constructs with VEGFR2 following their co-transfection in HEK 293T cells. HA immunoprecipitation is followed by western blotting for HA and VEGFR2. (E) Western blot analysis of VEGF induced VEGFR2 and ERK phosphorylation with and without FITC-L10 peptide (n=5). GAPDH was used as a loading control, with separate blots for total proteins (tp) and phosphorylated proteins (pp). *: P<0.05, **: P<0.01, ***: P<0.001, ****: P<0.0001. Statistical significance was determined by two-way ANOVA and Tukey’s multiple comparison post-hoc test.

A comparison of the rat, mouse, and human DCBLD2 amino acid sequences showed that while the three proteins share a poly-L segment in their N-terminal, the L5VL5 is replaced by L9 in murine and rat proteins (Fig. S13). We therefore generated a set of plasmids that express a 9 or 16 leucine peptide connected to a C-terminal GFP-HA tag. Following co-transfection with VEGFR2 in HEK 293 cells, co-immunoprecipitation studies showed that the removal of the valine in the L5VL5 sequence does not affect its interaction with VEGFR2. Interestingly however, elongating the polyleucine moiety to 16 amino acids, hampered VEGFR2 co-immunoprecipitation (Fig. 4C). Evaluation of additional constructs encompassing 7 to 12 leucine residues showed that the 9 to 11-leucine sequences appear to have the strongest interaction with VEGFR2, and this interaction diminishes as the number of leucine residues increases or decreases (Fig. 4D). Finally, to evaluate the functional significance of this interaction, the effect of a L10 synthetic peptide, FITC-L10EDAGA (connected to EDAGA to increase water solubility) on VEGF signaling was evaluated. Like FITC-traC, the FITC-L10 peptide enhanced VEGF signaling in HUVEC (Fig. 4E).

## DISCUSSION

This study uncovered a new non-canonical function for the unusually long signal sequence of DCBLD2. DCBLD2 regulates developmental and adult angiogenesis by engaging in a macromolecular complex with VEGFR2, which promotes VEGF signaling by dissociating VEGFR2 away from its negative regulators of downstream signaling, VE-cadherin and protein tyrosine phosphatases, PTP1 and TC-PTP ^13^. We identified the hydrophobic ‘traC’ subdomain of the DCBLD2 SS, and more specifically, its L5VL5 segment, as a key component of the DCBLD2 interaction with VEGFR2. The SS is not excised in the mature ESDN protein and potentially functions as a second transmembrane domain. Notably, the synthetic FITC-traC peptide specifically interacted with VEGFR2 and enhanced downstream signaling in a concentration-dependent manner, promoting VEGF-stimulated and ischemia-induced angiogenesis.

Signal sequences are typically composed of 25-30 amino acids. Many are cleaved off the nascent protein by signal peptidase I as signal peptides. The latter is often rapidly degraded by signal peptide peptidase after directing the protein to the endoplasmic reticulum ^22^. However, in some cases, the signal peptide remains stable after cleavage and may be involved in additional post-targeting functions as transmembrane or cytosolic/luminal endoplasmic reticulum peptides ^4, 23^. Non-cleaved signal sequences remain an integral component of the mature protein as signal anchor sequences ^24^. Several characteristics of the signal sequence, including its length, hydrophobicity, and charge of the residues surrounding the transmembrane core, influence the topology of a signal anchor ^25^. Long signal sequences (>40 residues) are less common in eucaryotes and are speculated to have additional post-targeting functions ^26^. Many of these long signal sequences may have a modular, bipartite ‘NtraC’ domain structure, with the C-domain performing as a fully functional SP for endoplasmic reticulum targeting ^5^. The N domain, on the other hand, may have additional targeting functions. In the case of Shrew-1 and DCBLD2, it is reported that the SS N domain may target the protein to mitochondria ^6, 24, 25^. Our data indicate that the DCBLD2 SS is not cleaved and therefore, may function as a signal anchor sequence. This would indicate that DCBLD2 is a double-pass transmembrane protein with cytosolic N and C terminals. Transmembrane domains are often engaged in protein-protein interactions and regulate the activity of the parent molecule. Similarly, the DCBLD2 SS and more specifically, its traC subdomain is involved in DCBLD2-VEGFR2 interaction and modulates VEGF signaling.

We further determined that the smallest unit in the DCBLD2 SS that can interact with VEGFR2 is the L5VL5 sequence. It is reported that most short ploy-leucine transmembrane domains containing single amino acid substitutions can bind to and activate PDGFRβ or the erythropoietin receptor ^27^. Minor changes in the composition of these sequences (e.g., changing the placement of a single side chain methyl group) can alter their partner selectivity ^28, 29^. Interestingly, even after removing the central valine, the L10 sequence could also interact with VEGFR2. However, shorter or longer poly-leucines were less effective in this regard, raising the possibility that the size of the poly-leucine sequence is one of the fundamental principles that determine such interactions. Leucine repeats are common in signal sequences, especially in higher organisms ^30^, and their conservation in mammals points to specific functions. An example of such unique functions could be the regulation of VEGFR2 signaling by the DCBLD2 SS L5VL5. While we have shown that the DCBLD2 SS shows some specificity with regards to its interactions with VEGFR2 (compared to PDGFRβ), the possibility of its involvement in other signaling pathways not tested here cannot be ruled out.

Notably, the exogenous FITC-traC and its derivative polyleucine peptide mimicked the effect of endogenous DCBLD2 in interacting with VEGFR2 and regulating VEGF signaling. This is probably related to the observation that FITC-traC is a cell-penetrating peptide and localizes within minutes in the cell membrane. It is reasonable to speculate that the latter location is where FITC-traC interacts with VEGFR2. These observations raised the possibility that FITC- traC and related peptides may serve as modulators of VEGFR2 signaling in vivo.

VEGFR2 is a main receptor for VEGF, the prototypic pro-angiogenesis growth factor, in endothelial cells ^31^. Angiogenesis is a key process during development, growth, and adulthood. Inadequate neovessel formation is a major pathologic feature in ischemia-related cardiovascular diseases and contributes to poor wound healing. In other pathological settings, e.g., diabetic retinopathy and malignancy, angiogenesis is improperly exaggerated. As such, there is considerable interest in regulators of VEGF signal transduction. FITC-traC promoted angiogenesis, not only when induced by exogenous VEGF, but also in the presence of ischemia, suggesting that it may be of value in treating ischemic diseases. Notably, FITC-traC also improved hindlimb blood flow recovery within a few days after femoral artery ligation in mice. This is in line with the finding that deletion of DCBLD2 impairs blood flow recovery in the same animal model ^13^, and may indicate that in addition to regulating angiogenesis, DCBLD2 and its SS derivatives may regulate collateral recruitment or arteriogenesis. The role of DCBLD2 SS in related signaling pathways and cell types will be the subject of future studies, which could also identify the specific VEGFR2 sequences/domains that interact with traC.

In conclusion, the DCBLD2 SS has a novel post-targeting function in interacting with and regulating VEGFR2 signaling and promoting angiogenesis through its transmembrane traC subdomain. This effect is narrowed down to the L5VL5 and L10 sequences. Therefore, besides their potential role as therapeutic targets ^32–34^, this study raises the prospect of another aspect of SS biology, i.e., their involvement in protein-protein interactions, which may be targeted by agonists (as shown here for DCBLD2 traC in promoting angiogenesis) or antagonists to develop therapeutic agents.

## Supplementary Material

Figures S1 to S13

Supplemental Statistical Analysis Data

Major Resources Table

## Non-standard Abbreviations and Acronyms

AKT: Protein kinase B
BSA: Bovine serum albumin
CD31: Cluster of differentiation 31
CD102: Cluster of differentiation 102
Cdh5: Cadherin 5 (VE-Cadherin)
DAPI: 4’,6-diamidino-2-phenylindole
DCBLD2: Discoidin, CUB, and LCCL domain containing 2
DMEM: Dulbecco’s modified eagle medium
EC: Endothelial cell
ECGS: Endothelial cell growth supplement
ER: Endoplasmic reticulum
ERK: Extracellular signal-regulated kinase
ESDN: Endothelial and smooth muscle cell-derived neuropilin-like protein
FBS: Fetal bovine serum
GFP: Green fluorescent protein
GAPDH: Glyceraldehyde 3-phosphate dehydrogenase
HA: Hemagglutinin
HBSS: Hank’s balanced salt solution
HEK 293T: Human embryonic kidney 293 cells transformed with SV40 large T antigen
HUVEC: Human umbilical vein endothelial cells
MLEC: Murine lung endothelial cells
mTOR: Mechanistic target of rapamycin
OCT: Optimal cutting temperature compound
PAD: Peripheral artery disease
PBS: Phosphate buffered saline
PDGFRβ: Platelet-derived growth factor receptor beta
PI3K: Phosphoinositide 3-kinase
poly-HEMA: Poly(hydroxyethyl methacrylate)
PTP1B: Protein tyrosine phosphatase 1B
SDS-PAGE: Sodium dodecyl sulfate polyacrylamide gel electrophoresis
SS: Signal sequences
SP: Signal peptide
TCPTP: T cell protein tyrosine phosphatase
VEGF: Vascular endothelial growth factor
VEGFR2: Vascular endothelial growth factor receptor 2

## Acknowledgments

We would like to thank the Yale West Campus Imaging Core for their support. We also extend our gratitude to Dr. Daniel DiMaio for sharing his expertise in biologically active simple peptides, Dr. Muhammad Riaz and Dr. Dongying Chen for providing technical assistance, and Dr. Michael Simons for his insightful comments on an earlier draft of the manuscript.

## Author contributions

Conceptualization: JJJ, MMS

Methodology: MG, LW, JJJ, ET, GK, AN, AAA, MZR, AG, KH, LN, JZ

Funding acquisition: MMS

Project administration: MMS

Supervision: MMS

Writing – original draft: MG, LW

Writing – review & editing: MG, LW, JJJ, ET, GK, AN, AAA, MZR, AG, KH, LN, JZ, MMS

## Data and materials availability

All data are available in the main text or the supplementary materials.

## What is Known?

- The Discoidin, CUB, and LCCL domain containing 2 (DCBLD2) protein is a type I transmembrane protein with a long signal sequence that interacts with VEGFR2 and modulates angiogenesis.
- Signal sequences typically guide proteins to the endoplasmic reticulum. They may be cleaved as signal peptides which are often degraded after their targeting function.
- Long signal sequences are rare and have been speculated to serve additional functions, but their precise roles are unknown.

## What New Information Does This Article Contribute?

- The DCBLD2 signal sequence is not cleaved from the mature protein and contains a hydrophobic transmembrane subdomain (traC) that interacts with VEGFR2. Synthetic traC peptide promotes VEGF signaling and angiogenesis.
- The L5VL5 sequence within the traC subdomain is crucial for VEGFR2 interaction, with synthetic peptides mimicking this sequence enhancing VEGF signaling.
- Long signal sequences may play roles in protein-protein interactions and cell signaling.

## Summary of Novelty and Significance

We identified a novel function for the atypically long signal sequence of DCBLD2 in regulating VEGFR2 signaling and angiogenesis. The DCBLD2 signal sequence remains uncleaved in mature protein and has a hydrophobic subdomain, traC, that interacts directly with VEGFR2.

The L5VL5 sequence within traC is involved in this interaction, and synthetic peptide containing traC and related sequences enhance VEGF signaling and promote angiogenesis. These findings not only underscore the potential of the DCBLD2 signal sequence and its derivatives as therapeutic agents in conditions where angiogenesis is impaired, but also prompt further exploration into the broader roles of long signal sequences in vascular biology, development, and disease.

